# Investigating resting-state functional connectivity in the cervical spinal cord at 3T

**DOI:** 10.1101/073569

**Authors:** Falk Eippert, Yazhuo Kong, Anderson M. Winkler, Jesper L. Andersson, Jürgen Finsterbusch, Christian Büchel, Jonathan C.W. Brooks, Irene Tracey

## Abstract

The study of spontaneous fluctuations in the blood-oxygen-level-dependent (BOLD) signal has recently been extended from the brain to the spinal cord. Two ultra-high field functional magnetic resonance imaging (fMRI) studies in humans have provided evidence for reproducible resting-state connectivity between the dorsal horns as well as between the ventral horns, and a study in non-human primates has shown that these resting-state signals are impacted by spinal cord injury. As these studies were carried out at ultra-high field strengths using region-of-interest (ROI) based analyses, we investigated whether such resting-state signals could also be observed at the clinically more prevalent field strength of 3T. In a reanalysis of a sample of 20 healthy human participants who underwent a resting-state fMRI acquisition of the cervical spinal cord, we were able to observe significant dorsal horn connectivity as well as ventral horn connectivity, but no consistent effects for connectivity between dorsal and ventral horns, thus replicating the human 7T results. These effects were not only observable when averaging along the acquired length of the spinal cord, but also when we examined each of the acquired spinal segments separately, which showed similar patterns of connectivity. Finally, we investigated the robustness of these resting-state signals against variations in the analysis pipeline by varying the type of ROI creation, temporal filtering, nuisance regression and connectivity metric. We observed that – apart from the effects of band-pass filtering – ventral horn connectivity showed excellent robustness, whereas dorsal horn connectivity showed moderate robustness. Together, our results provide evidence that spinal cord resting-state connectivity is a robust and spatially consistent phenomenon that could be a valuable tool for investigating the effects of pathology, disease progression, and treatment response in neurological conditions with a spinal component, such as spinal cord injury.

## Introduction

The temporal and spatial organization of intrinsic brain activity is currently a subject of intense research. Functional magnetic resonance imaging (fMRI) studies have shown that spontaneous fluctuations in the blood-oxygen-level-dependent (BOLD) signal are organized into distinct and reproducible resting-state networks, such as the sensorimotor, default-mode, or executive-control networks (Buckner et al., 2013; Fox and Raichle, 2007; Power et al., 2014). With the neurophysiological origin of these resting-state signals becoming more evident (Leopold and Maier, 2012; Schölvinck et al., 2013) and their clinical relevance more appreciated (Fox and Greicius, 2010; Zhang and Raichle, 2010), they are increasingly used to probe the integrity and properties of neural circuits in health and disease.

These organized resting-state fluctuations are not an exclusively cortical phenomenon, but have also been observed in subcortical regions as low as the pons and medulla (Beissner et al., 2014; Bianciardi et al., 2016), raising the question whether they constitute a functional signature of the entire central nervous system and might thus be detectable in the spinal cord as well. However, answering this question is a difficult endeavour, because it is challenging to obtain reliable fMRI data from the spinal cord due to a number of issues (Giove et al., 2004; Stroman et al., 2014; Summers et al., 2010), the most prominent of which are: 1) the spinal cord has a very small cross-sectional area (Fradet et al., 2014; Ko et al., 2004), 2) the detrimental influence of physiological noise from cardiac and respiratory sources is much more prominent in the spinal cord than in the brain (Piché et al., 2009; Verma and Cohen-Adad, 2014), and 3) signal loss and image distortion periodically occur along the spinal cord due to the different magnetic susceptibility of vertebrae and connective tissue (Cooke et al., 2004; Finsterbusch et al., 2012).

Despite these obstacles, a few groups have started investigating spinal cord resting-state functional connectivity. Wei and colleagues (2010), for instance explored resting-state signals in the human spinal cord by using independent component analysis (ICA), and reported that the networks detected at the single-subject level were dominated by signal in the frequency range of the respiratory cycle – thus hindering an unequivocal interpretation with regard to a neuronal origin. Building on this initial finding, two other exploratory ICA-based studies used comprehensive denoising strategies and group-level analyses to demonstrate spatially distinct and reproducible spinal cord resting-state signals that are likely to be of neuronal origin (Kong et al., 2014; San Emeterio Nateras et al., 2016). Barry and colleagues used ultra high field imaging at 7T in combination with a hypothesis-driven region-of-interest (ROI) approach to demonstrate robust and reproducible resting-state functional connectivity in the human spinal cord (Barry et al., 2014, 2016). They observed significant time-course correlations between the ventral horns as well as between the dorsal horns at the group level, but not between ventral and dorsal horns. The clinical significance of such resting-state connectivity was recently demonstrated in a non-human primate model of spinal cord injury, where a spatially-specific influence of lesions on spinal cord functional connectivity was observed (Chen et al., 2015).

These studies hint at the translational potential of using spinal cord resting-state fMRI signals in clinical situations that involve spinal pathology and where a non-invasive metric of disease progression and treatment response would be welcome; examples include multiple sclerosis, spinal cord compression, spinal cord injury, and chronic pain (Wheeler-Kingshott et al., 2014). However, if resting-state connectivity of the human spinal cord is indeed to be used as a potential biomarker for disease progression or treatment effects, some conditions need to be satisfied. First, we need to be able to successfully acquire these signals at the clinically relevant field strength of 3T, because 7T scanners are currently only available in a small minority of research-focussed departments (less than 50 worldwide; Balchandani and Naidich, 2015). Second, we need to demonstrate that the technique is able to show robust results when obtaining data from distinct spinal segments, because spinal pathologies can be very localised (for example in spinal cord compression; Nouri et al., 2015). Finally, we need to show that the results we obtain are robust against variations in the data analysis pipeline, thus ensuring their inferential reproducibility (Goodman et al., 2016). We consider these three conditions as preliminary but important steps with respect to feasibility, which need to occur before setting sight on longer term goals, such as carrying out longitudinal studies on the stability of spinal cord resting-state connectivity and formal sensitivity/specificity analyses in patient cohorts.

Here, we evaluated to what extent spinal cord resting-state connectivity can satisfy the above-mentioned conditions by reanalysing a previously published data-set that was acquired at a field strength of 3T and used ICA to explore spinal cord resting-state signals (Kong et al., 2014). First, we used this data-set to test whether we could replicate the previously obtained 7T results (significant connectivity between ventral horns and between dorsal horns after averaging over several segments; Barry et al., 2014). Next, we tested whether these results also held for distinct spinal segments - an approach that has only now become possible with the development of a probabilistic atlas of spinal cord segments in a standard space (Cadotte et al., 2015; De Leener et al., 2016). Third, we assessed whether the obtained results were stable across variations of our data analysis pipeline, by varying the type of 1) ROI creation, 2) temporal filtering, 3) nuisance regression, and 4) connectivity metric. Together, these tests should allow us to determine whether resting-state connectivity in the human spinal cord might be a useful tool in both basic neuroscience and clinical investigations.

## Methods

### Participants

This study is based on a re-analysis of the data presented in Kong et al. (2014) and thus contains data from the same 20 healthy male participants (age: 26.5 ±3.9 years). The Ethics Committee of the Medical Board in Hamburg, Germany, approved the study and all participants gave written informed consent.

### Data acquisition

Magnetic resonance imaging (MRI) data were acquired in an eyes-open state on a 3T system (Magnetom Trio, Siemens, Erlangen, Germany). fMRI data were collected as the last session in a larger spinal fMRI experiment consisting of two sensory and two motor sessions using a recently developed slice-specific z-shim protocol (Finsterbusch et al., 2012). During all sessions a white crosshair was shown on the screen, which turned red every 15s; participants were asked to stay as still as possible and movements were limited by using a vacuum cushion. Participants were imaged with a 12-channel head coil combined with a 4-channel neck coil (both receive-only), with the cervical spinal cord centred in the neck coil and positioned at isocenter in the magnet. Functional images were acquired using a T2*-weighted gradient-echo echo-planar imaging (EPI) sequence (repetition time 1890ms, echo time 44ms, flip angle: 80°, field of view: 128x128mm², matrix: 128x128, GRAPPA with a PAT-factor of 2). We acquired 16 transversal slices using a slice thickness of 5mm in order to achieve an adequate signal-to-noise ratio despite our high in-plane resolution (1x1mm²). The resulting target volume covered the spinal cord from the 4^th^ cervical vertebra to the 1^st^ thoracic vertebra – based on probabilistic maps of spinal levels, this volume includes segments C6, C7, C8, and T1 (Cadotte et al., 2015). To minimize sensitivity to flow effects, first-order flow compensation in the slice direction of both the slice-selection and the z-shim gradient pulse and spatially-selective saturation pulse superior and inferior to the target volume were used and the images obtained with the individual coil channels were combined with a sum-of-squares algorithm. Furthermore, additional saturation pulses were applied posterior and anterior to the target region, i.e. in the phase-encoding direction, in order to avoid pulsatile blood flow artefacts. Periodic signal dropout due to magnetic field inhomogeneity induced by the alternation of vertebrae and connective tissue was minimized by using slice-specific z-shimming, which has been shown to lead to a reduction in signal intensity variation along the cord of ~80% (Finsterbusch et al., 2012). The adjustments prior to the functional acquisitions (i.e. shimming) were performed on a manually defined volume of about 35x30x70mm³ covering the target region in the spinal cord. Only the neck-coil was used for acquisition of fMRI data and a total of 250 volumes were acquired for each participant (7.5-minute scanning time). Please note that three initial volumes (occurring before the 250 volumes used for analysis) were used to achieve steady-state conditions and to acquire reference data for GRAPPA; these volumes were thus not included in the resting-state data analysis. For the employed repetition time and flip angle this means that all included signals were within less than 0.005% and 0.2% of the steady-state signal for tissue and cerebrospinal fluid, respectively, which is one to three orders of magnitude lower than the noise level. To monitor cardiac and respiratory signals during fMRI data acquisition, participants wore a pulse oximeter and respiratory belt, and physiological data were recorded together with the trigger pulses preceding the acquisition of each volume.

We also acquired high-resolution (1x1x1mm³) T_1_-weighted anatomical images using a 3D-MPRAGE sequence (sagittal slice orientation, repetition time 2.3s, echo time 3.5ms, flip angle 9^°^, inversion time 1.1s, field-of-view 192x240x256mm³). The field of view for this acquisition covered an area that spanned at least from the midbrain to the second thoracic vertebra in every participant; both the neck coil and the head coil were used for this acquisition.

### Data processing

Data were processed using tools from FSL (FMRIB Software Library; http://fsl.fmrib.ox.ac.uk/fsl/fslwiki/; Jenkinson et al., 2012). First, each slice was motion corrected for x- and y-translations using FLIRT (FMRIB’s Linear Image Registration Tool; http://fsl.fmrib.ox.ac.uk/fsl/fslwiki/FLIRT; Jenkinson et al., 2002); translations in the z-direction and rotations were assumed to be minimal, which was confirmed by visual inspection (focussed on the intervertebral disks for translations in the z-direction and on the spinal cord for rotations) following motion correction. Note that slice-wise motion correction can outperform volume-wise approaches, because spinal cord displacement varies along the rostro-caudal axis of the spinal cord according to the cardiac (Figley and Stroman, 2007) and respiratory cycle (Verma and Cohen-Adad, 2014).

Next, we used FEAT (FMRI Expert Analysis Tool; http://fsl.fmrib.ox.ac.uk/fsl/fslwiki/FEAT) to carry out physiological noise regression and high pass filtering (using a cut-off of 100s) – these two steps were performed simultaneously in order to avoid spectral misspecification (Hallquist et al., 2013). The influence of physiological noise of cardiac and respiratory nature is particularly pronounced in the spinal cord and we thus used PNM (Physiological Noise Modelling; http://fsl.fmrib.ox.ac.uk/fsl/fslwiki/PNM; Brooks et al., 2008) in the context of FEAT to remove these noise sources. PNM is based on the RETROICOR approach (Glover et al., 2000) and removes physiological confounds from motion-corrected data using slice-specific regressors based on the calculated phase for each slice relative to the cardiac and respiratory cycles (see also Kong et al., 2012). In the physiological noise model we used here, cardiac, respiratory and interaction effects were modelled using Fourier series, resulting in a total of 32 regressors. Additional nuisance regressors consisted of a) low frequency cerebrospinal fluid (CSF) signal (extracted from voxels whose variance lay in the top 10 percentile within a region including both the spinal cord and CSF space), b) heart rate (value of the smoothed beats per minute (BPM) trace at the acquisition time for each slice), c) motion correction parameters (x and y translation), and d) a regressor that modeled the colour-change of the cross-hair that was presented on the screen. The obtained residuals from each fMRI scan (i.e. physiological noise corrected and high-pass filtered data) were used for further analysis.

Finally, we brought the residuals of the functional data into a common anatomical space. The registration of functional images to the structural volume was initialised using the scanner sqform transformation. Due to EPI distortion in the fMRI data,there remained residual mismatch between the structural and functional data in some slices following the initial transformation. We therefore applied an additional slice-wise registration procedure (x and y translations) on these data (based on hand-drawn spinal cord masks), which minimised the mismatch, and brought fMRI data into good alignment with each participant’s structural volume. We then identifieda participant with minimal anterior-posterior and left-right curvature of the spine,which became the experimental template. Subsequently, each participant’s structural image was registered to this template using a two-step procedure: we first used FLIRT (FMRIB’s Linear Image Registration Tool; http://fsl.fmrib.ox.ac.uk/fsl/fslwiki/FLIRT; Jenkinson and Smith, 2001) with default options and angular search range set to 0 degrees and then employed the resulting transformation matrix as a starting point for FNIRT (FMRIB’s Non-Linear Image Registration Tool; http://fsl.fmrib.ox.ac.uk/fsl/fslwiki/FNIRT; Andersson et al., 2010); important details include: 40 iterations, changing warp resolution from 10mm to 1mm, bias field modelling resolution of 20mm and weighting lambda of 100, weighting mask covering spinal cord and disks/vertebrae. The sqform, XY translation and non-linear warping transformations were then applied to the residuals of the functional data to bring them into a common anatomical space (resampled at 1x1x1mm).

#### Data analysis - Aim 1

In order to test our first aim - whether we could replicate the results of the recent ROI-based 7T resting-state reports (Barry et al., 2016, 2014) at our field strength of 3T - we needed to create masks for each of the four grey matter horns (along the length of the spinal cord, with our field of view covering segments C6 to T1). These masks were based on a probabilistic grey matter atlas (Taso et al., 2014) that is integrated with the MNI-Poly-AMU template of the spinal cord (Fonov et al., 2014) and is available within SCT (Spinal Cord Toolbox; https://sourceforge.net/projects/spinalcordtoolbox; De Leener et al., 2016). We obtained these four masks by 1) thresholding the probabilistic grey matter atlas at 50%, 2) splitting the supra-threshold image into 4 different images (one for each horn), 3) making sure that there was at least a one-voxel gap between dorsal and ventral horns, and 4) making sure that the minimal distance between the ventral horns was equal to the minimal distance between the dorsal horns (which resulted in discarding grey matter voxels that belonged to the central grey matter) so that a different distance would not bias the correlation; all steps were done separately for each slice and at the end all slices were merged together. For time-course extraction and statistical analysis, please see below.

#### Data analysis - Aim 2

In order to test our second aim - whether spinal cord resting-state connectivity could be observed at single segments at our field strength of 3T - we created masks for each horn that did not span the whole extent of the cord, but were instead limited to single spinal cord segments (C6, C7, C8, and T1 in our case). These masks were created by intersecting the previously obtained horn-specific masks with probabilistic masks defining spinal cord segments (Cadotte et al., 2015), which are also integrated with the MNI-Poly-AMU template and available within SCT. We thresholded the probabilistic spinal segment masks at 50% and minimally edited them manually (removing any overlap between neighbouring segments).

In addition to creating the masks needed to assess Aim 1 (whole-cord correlations) and Aim 2 (single-segment correlations), we also needed to obtain a mapping between our common anatomical space (in which our normalized structural and functional images reside) and the space in which the probabilistic spinal cord atlases reside and where the masks were defined. In order to do so, we first averaged our individual normalized structural images and then applied a non-rigid registration procedure to the resulting average normalized structural image, using the MNI-Poly-AMU template as a target. This was done using procedures implemented in SCT (for details, see De Leener et al., 2014; Fonov et al., 2014) and resulted in deformation fields describing the mapping between the two spaces, allowing us to bring the masks into our common anatomical space.

For both Aim 1 and Aim 2 (where all the following steps were carried out per spinal segment), we used the following procedures to estimate resting-state connectivity. We 1) obtained the average time-course from each of the four horn masks in each participant, 2) calculated Pearson’s correlation coefficients between the time-courses for all four horn masks in each participant, 3) averaged the correlation coefficients from left-dorsal-with-left-ventral and right-dorsal-with-right-ventral correlations (to create an index for within-hemicord dorsal-ventral correlations) as well as left-dorsal-with-right-ventral and right-dorsal-with-left-ventral correlations (to create an index for between-hemicord dorsal-ventral correlations) in each participant, and 4) used non-parametric permutation tests for group-level inference. With regard to this last point, we assessed whether the average across subjects of each of the four horn-to-horn correlations (1: dorsal-dorsal, 2: ventral-ventral, 3: within-hemicord dorsal-ventral, 4: between-hemicord dorsal-ventral) was different than zero using permutation testing as implemented in PALM (Permutation Analysis of Linear Models, http://fsl.fmrib.ox.ac.uk/fsl/fslwiki/PALM; Winkler et al., 2014); we used 10000 sign-flips for each test and report two-tailed family-wise-error (FWE) corrected p-values (adjusted for the four tests performed). To give an insight into the inter-individual variability of the four horn-to-horn correlations, we also report the percentage of participants who show each effect. For the whole-cord analysis (Aim 1) we also compared the strength of the different horn-to-horn correlations (1: dorsal-dorsal vs ventral-ventral, 2: dorsal-dorsal vs within-hemicord dorsal-ventral, 3: dorsal-dorsal vs between-hemicord dorsal-ventral, 4: ventral-ventral vs within-hemicord dorsal-ventral, 5: ventral-ventral vs between-hemicord dorsal-ventral, 6: within-hemicord dorsal-ventral vs between-hemicord dorsal-ventral) using the permutation testing analogue of a paired t-test implemented in PALM; we used 10000 permutations and again report two-tailed FWE-corrected p-values (adjusted for the six tests performed). Please note that we report group-averaged correlation coefficients, which tend to exhibit a small conservative bias, i.e. will slightly underestimate the true correlation (compared to averaging Fisher-z transformed correlation coefficients and then back-transforming the average to Pearson’s r, which will slightly overestimate the true correlation; Clayton and Dunlap, 1987; Corey et al., 1998).

#### Data analysis - Aim 3

In order to test our third aim - which was to assess how robust the observed resting-state connectivity would be against variations in the analysis pipeline, i.e. how reproducible / robust the results would be - we varied the type of 1) ROI creation, 2) temporal filtering, 3) nuisance regression, and 4) connectivity metric.

- *ROI creation*: We not only used the above-described probabilistic masks for each horn (which we will refer to as PROB [for probabilistic]), but also created masks that consisted of just one voxel for each horn (per slice), located at the x-y centre of gravity of each horn (which we will refer to as COG [for centre of gravity]); please note that while the x-y centre of gravity was calculated per slice, we then combined these voxels across all slices to create the COG masks. These masks were created to assess the effects that differences in ROI creation can have on connectivity estimates (Marrelec and Fransson, 2011; Smith et al., 2011) and to ameliorate several issues that could potentially complicate interpreting the data when using the PROB masks: 1) different number of voxels in ventral horns vs dorsal horns (with COG, we just have one voxel per slice per horn), 2) influence of residual CSF fluctuations (with COG, the masks are further away from the subarachnoid space), 3) influence of signal from large veins at the edge of the cord (Cohen-Adad et al., 2010; with COG, the masks are further away from the cord edge), and 4) signal overlap between dorsal horns and ventral horns (due to the point-spread-function of the BOLD response and there only being a one-voxel gap between dorsal horn and ventral horn masks; with COG, the dorsal and ventral horn masks are more strongly separated from each other).
- *Temporal filtering*: Resting-state data have traditionally been band-pass filtered due to the assumption that connectivity is driven by low-frequency fluctuations. However, this has recently been challenged with the discovery that high frequencies also contain meaningful signal (Chen and Glover, 2015; Niazy et al., 2011). At 7T, Barry and colleagues (Barry et al., 2016) showed that this also holds true for the spinal cord when they noticed that frequencies above 0.08Hz carried meaningful signal, as for example evidenced by higher reproducibility of spinal cord resting-state correlations across sessions. We therefore evaluated the effects of using only a high-pass filter (with a cut-off of 100s, i.e. 0.01Hz; which we will refer to as HP) or using a band-pass filter with a pass-band between 0.01 and 0.08Hz (similar to Barry et al., 2014; which we will refer to as BP). Note that when only using a high-pass filter, we can obtain signals up to the Nyquist frequency of 0.26Hz.
- *Nuisance regression*: We investigated several different slice-wise nuisance regression options in addition to the previously applied slice-wise PNM (see section “Data processing”). First, we investigated the effect of regressing out the average white matter (WM) signal per slice, which could help to mitigate residual physiological noise effects as well as time-dependent partial volume effects at the grey matter to white matter boundary due to residual motion (Barry et al., 2014). The white matter signal time-course was obtained from the probabilistic white matter mask (thresholded at 10%) - to minimize partial volume effects with grey matter, we subtracted a dilated version of the probabilistic grey matter mask (thresholded at 50%) from this mask. Second, we investigated the effect of regressing out residual cerebrospinal fluid (CSF) signals - in our original PNM, we used one regressor per slice to capture CSF signals, but this might not be sufficient due to CSF flow not being homogenous within the subarachnoid space (CSF flow occurs in different channels with different time-profiles; Schroth and Klose, 1992a; Henry-Feugeas et al., 2000). We therefore carried out a principal component analysis (PCA) on voxel-wise CSF time-courses (which we obtained from the CSF mask that is part of the MNI-Poly-AMU template) and used the first four principal components per slice as regressors (since there are four different CSF channels); together these 4 components explained almost 50% of the variance. Third, we investigated the effect of regressing out non-spinal (NS) signals, i.e. signals that are clearly non-neuronal in origin (e.g. signals in connective tissue or muscles, remaining vascular signals, wide-spread intensity fluctuations due to swallowing, image artefacts, etc.) but might impact on spinal cord BOLD fluctuations. We therefore carried out a PCA on voxel-wise NS time-courses (which we obtained by 1) combining the MNI-Poly-AMU cord mask and CSF mask, 2) dilating the resulting mask and 3) logically inverting the resulting mask) and used the first ten principal components per slice as regressors (each of which explained at least 1% of the variance). This resulted in a total of 8 possible nuisance regression combinations (1: none, 2: WM, 3: WM+CSF, 4: WM+NS, 5: WM+CSF+NS, 6: CSF, 7: CSF+NS, 8: NS).
- *Connectivity metric*: We not only used Pearson’s correlation coefficient as described above (which we will refer to as FULL, for full correlation, and which was used by Barry et al., 2014), but also used partial correlation and a regularized version of partial correlation. Partial correlation (which we will refer to as PARTIAL and which was used by Barry et al., 2016) estimates the correlation between two ROIs while controlling for the influence of the time-courses in the remaining two ROIs that do not enter the correlation, i.e. when assessing dorsal-dorsal connectivity this controls for any contributions from the ventral ROIs. This is an attractive approach that is not only able to distinguish between direct and indirect connections (Marrelec et al., 2006), but should also remove any remaining global signal fluctuations that are shared between the ROIs (e.g. residual movement or physiological noise effects).Regularized partial correlation (which we will refer to as REGPARTIAL)imposes a sparseness constraint on the partial correlation matrix and can be beneficial in situations where there are high noise levels, resting-state data have a short duration, or networks have a large number of ROIs. We used theFSLNETS (http://fsl.fmrib.ox.ac.uk/fsl/fslwiki/FSLNets) implementation ofregularized partial correlation (which is based on L1-norm regularization, see http://www.cs.ubc.ca/~schmidtm/Software/L1precision.html) with a regularisation-controlling parameter λ of 5 (Smith et al., 2011).

Combining the different factors of ROI creation (two), temporal filtering (two), nuisance regression (eight), and connectivity metric (three) resulted in a total of 96 analyses; for brevity, we only report the results from the whole-cord analyses. In order to gauge the robustness of each of the four horn-to-horn correlations (1: dorsal-dorsal, 2: ventral-ventral, 3: within-hemicord dorsal-ventral, 4: between-hemicord dorsal-ventral) against variations in data analysis, we first investigated whether the sign of the correlation changed with analysis choice, i.e. for each horn-to-horn correlation we report the number of positive correlations among all 96 performed analyses. After observing that only two of the four horn-to-horn correlations were robust against variations in data analysis, we then assessed how the significance of these correlations (again we use FWE-corrected two-tailed p-values as detailed above) was influenced by variations in data analysis, i.e. we report the number of significant correlations among all 96 performed analyses. Supplementing these descriptive reports of the binned data (i.e. positive / negative, significant / not significant), we used a four-way repeated measures analysis of variance (ANOVA; factors ROI creation [2 levels], temporal filtering [2 levels], nuisance regression [8 levels], and connectivity metric [3 levels]) for each of the four horn-to-horn correlation coefficients to investigate the impact of each factor.

Finally, we carried out two complementary analyses that made use of all the 96 different analyses in order to provide evidence for the existence of horn-to-horn connectivity that is identifiable at 3T. In a first analysis we averaged the correlation coefficients across the 96 analyses within each subject and horn-to-horn correlation and then carried out the same permutation test as mentioned above on these averages (again reporting two-tailed FWE-corrected p-values). In a second analysis we used the recently developed modification of non-parametric combination testing (NPC; Winkler et al., 2016) in order to perform joint inference across all 96 analyses using the Fisher combining function. As in the first analysis we report FWE-corrected p-values. Note that these two tests are not equivalent: the null hypothesis for the first test is that the average effect is zero, whereas the null hypothesis for the second test is that all the effects are zero.

## Results

### Control analyses

We first investigated whether the temporal signal-to-noise ratio (tSNR) would exhibit any systematic differences between the different horns. Group-averaged tSNR maps and estimates were obtained after motion correction, physiological noise modelling, high-pass filtering and registration to standard space; note that no smoothing was performed. As can be seen in Figure 1, the group-averaged tSNR i) was rather homogeneous within each horn (with a slight gradient of decreased tSNR towards the dorsal and ventral edges of the horn masks), ii) was nearly identical for the different horns within a segment (and showed small variation across participants, as evidenced by the small error bars), and iii) was very similar for segments C6 to C8, with a slight drop in segment T1.

**Figure 1.**
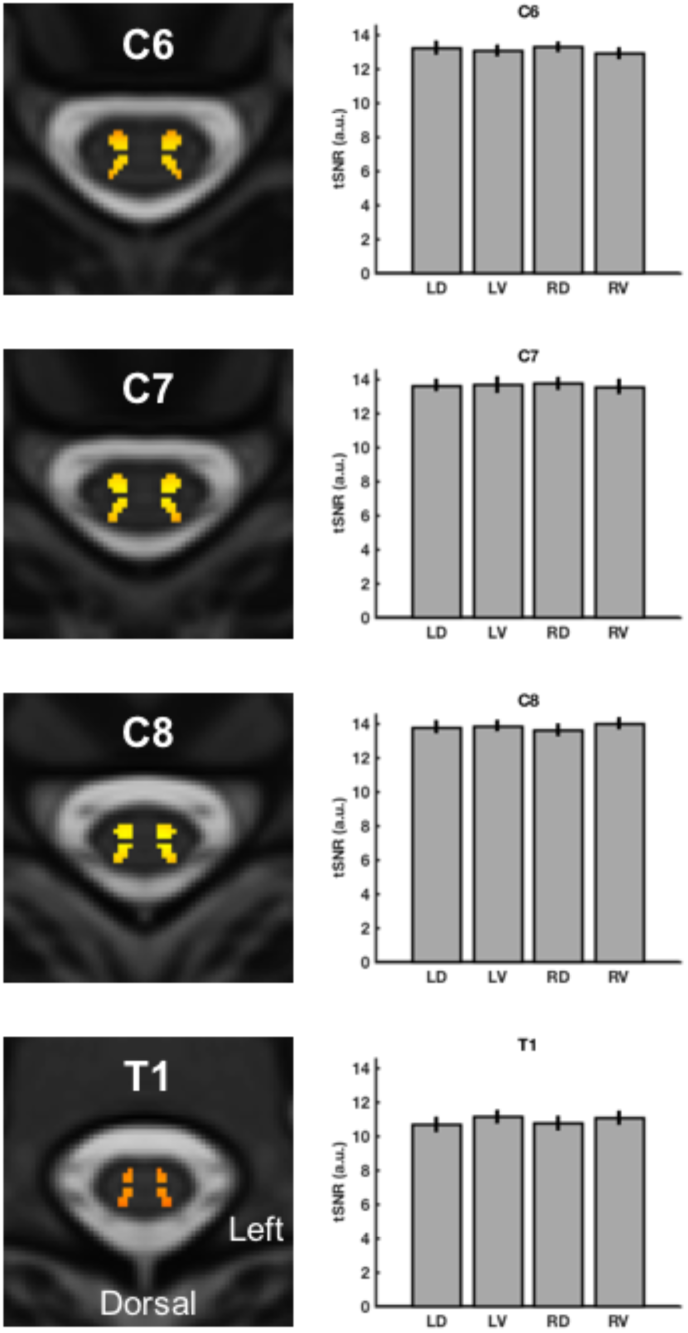
Group-averaged tSNR.

On the left side, group-averaged voxel-wise maps of the tSNR in the different spinal segments are displayed (transversal slice at the middle of each segment). The background image is the T2-weighted MNI-Poly-AMU template, the red-to-yellow coded tSNR is displayed only in voxels belonging to the probabilistic grey matter masks, and the colour scale is identical for all images. On the right, group-averaged segmental tSNR estimates are displayed for each horn (averaged across all voxels within a horn for each participant, i.e. taking into account all slices belonging to a segment). Both the maps on the left and the averaged estimates on the right are based on tSNR after motion correction, physiological noise modelling, and high-pass filtering (but no smoothing). Error bars represent the standard error of the mean. Abbreviations: LD, left dorsal horn; LV left ventral horn; RD, right dorsal horn; RV, right ventral horn.

In all subsequent analyses, we report one value for within-hemicord dorsal-ventral correlations (based on averaging correlation coefficients for “left-dorsal-with-left-ventral” and “right-dorsal-with-right-ventral”) and one value for between-hemicord dorsal-ventral correlations (based on averaging correlation coefficients for “left-dorsal-with-right-ventral” and “right-dorsal-with-left-ventral”). As this rests on the assumption that there are no meaningful laterality differences between the to-be-averaged correlations, we tested for this using two-tailed non-parametric permutation tests. We did not observe any significant laterality differences, neither for the within-hemicord dorsal-ventral correlations nor for the between-hemicord dorsal-ventral correlations. While one test showed a marginally significant result (whole-cord between-hemicord dorsal-ventral correlation: p = 0.05), this was far from significance when considering correction for multiple comparisons (p = 0.33). Figure 2 shows descriptively that there is no systematic laterality effect, with median values clustering around zero, both for the whole cord and the different segments.

**Figure 2.**
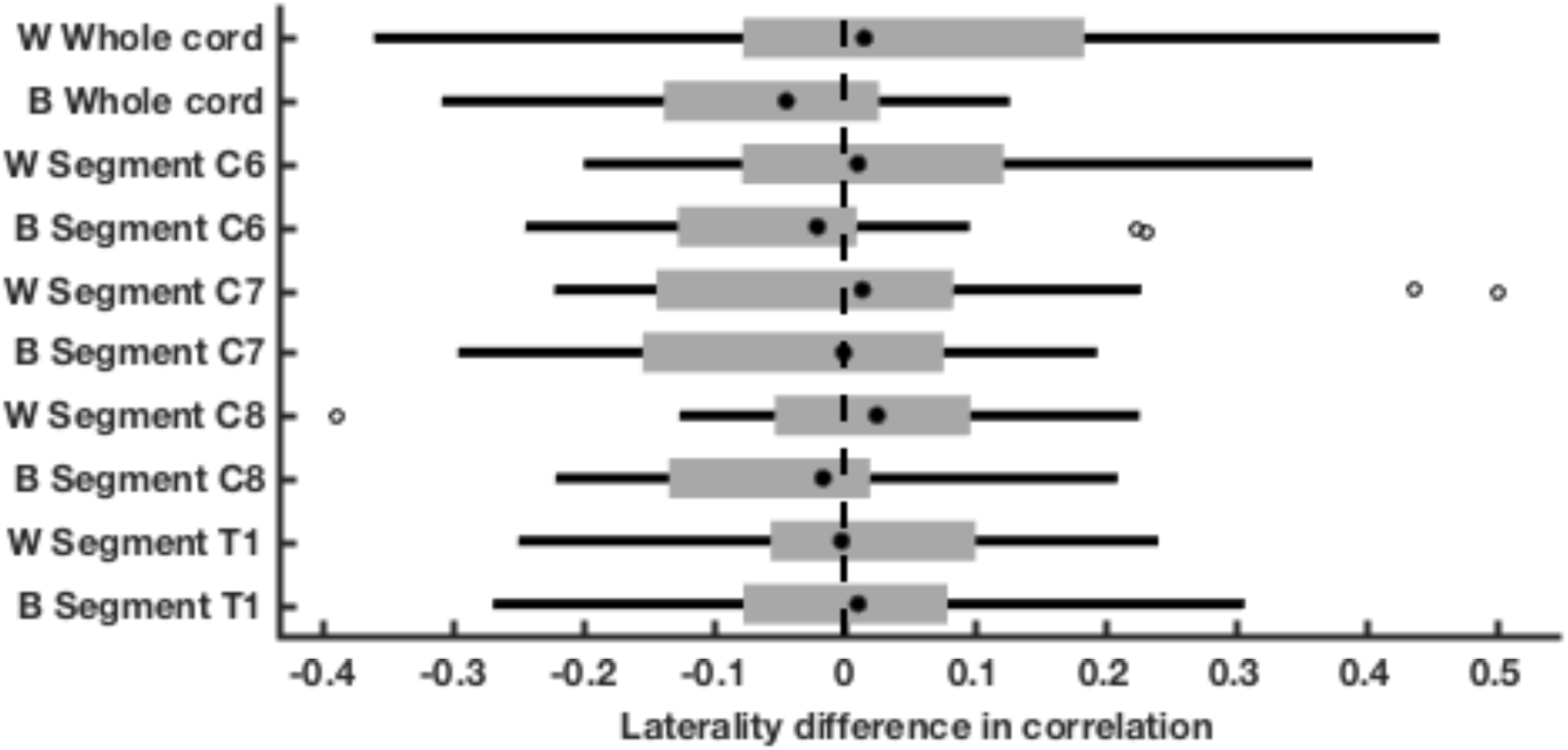
Laterality difference in dorsal-ventral correlations.

Both for the whole cord as well as for each segmental level, we calculated the laterality difference in dorsal-ventral horn correlation coefficients. The filled black dot represents the median, the edges of the boxes cover represent the 25^th^ and 75^th^ percentiles and whiskers encompass approximately 99% of the data; outliers are represented by non-filled circles. Abbreviations: W, within-hemicord dorsal-ventral correlations (positive difference reflects left-dorsal-with-left-ventral > right-dorsal-with-right-ventral); B between-hemicord dorsal ventral correlations (positive difference reflects right-ventral-with-left-dorsal > right-dorsal-with-left-ventral).

### Aim 1 - whole cord connectivity

Our first aim was to test whether we could replicate the results of the recent ROI-based 7T resting-state reports (Barry et al., 2016, 2014), namely significant time-course correlations between the two dorsal horns, as well as between the two ventral horns (but not between dorsal and ventral horns) when averaged over the acquired rostro-caudal extent of the spinal cord (Figure 3). Indeed, we observed that the dorsal horns exhibited significant functional connectivity (r = 0.22, p < 0.001), as did the ventral horns (r = 0.22, p < 0.001), with 90% of participants showing positive correlations between dorsal horns and 95% of participants showing positive correlations between ventral horns. In contrast to these robust findings, dorsal-ventral horn connectivity within a hemi-cord was just minimally above zero (r = 0.02, p = 0.93; 50% of participants showed positive correlations), whereas dorsal-ventral horn connectivity between hemicords was significantly negative (r = -0.08, p = 0.03; 30% of participants showed positive correlations; but see results described under Aim 3).

**Figure 3.**
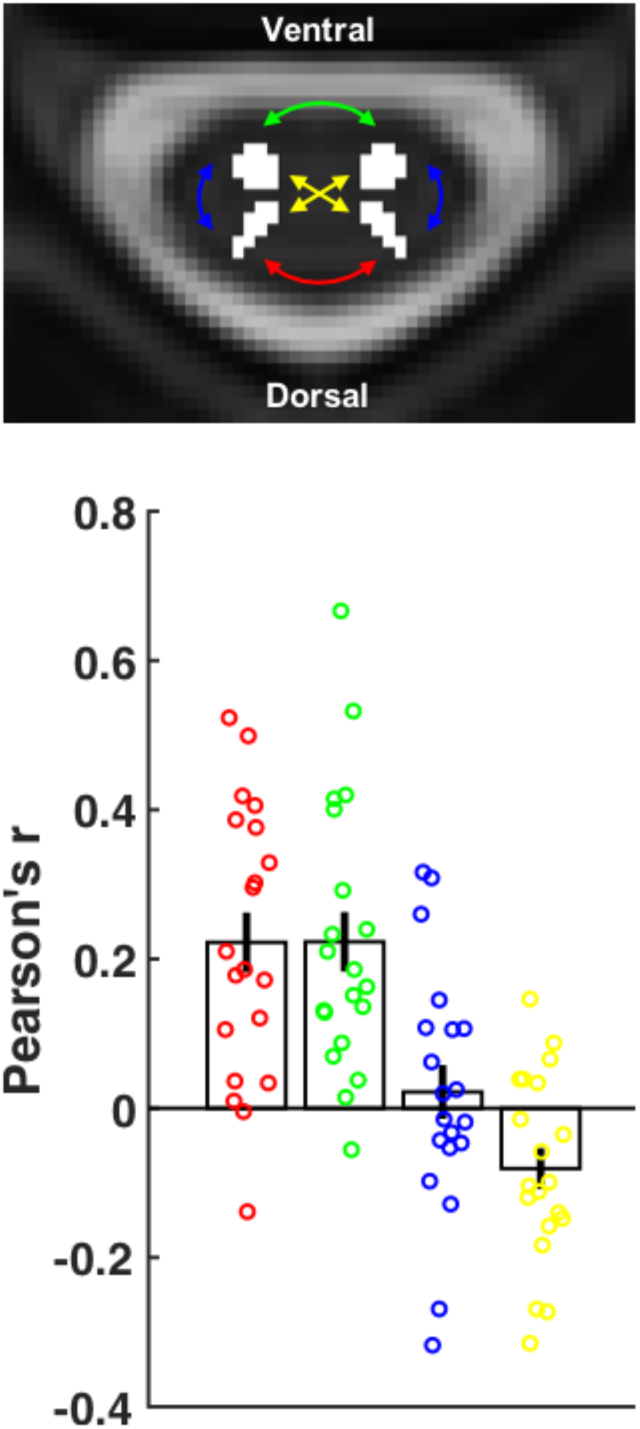
Connectivity averaged along the cord.

The transversal slice is taken from the T2-weighted MNI-Poly-AMU template at the middle of segment C6, with the four horn masks overlaid in white and coloured arrows indicating the four different types of horn-to-horn connectivity we investigated (dorsal-dorsal connectivity is depicted in red, ventral-ventral connectivity in green, within-hemicord dorsal-ventral connectivity in blue, and between-hemicord dorsal-ventral connectivity in yellow). The bar-plot displays the group averaged correlation (+/- the standard error of the mean) for each of the four horn-to-horn correlations and the circles indicate participant-specific correlations.

As an aside, we also compared the four different horn-to-horn correlations to each other. We observed that dorsal horn connectivity and ventral horn connectivity were both significantly stronger than a) within-hemicord dorsal-ventral horn connectivity (dorsal: p = 0.02, ventral: p = 0.02) and b) between-hemicord dorsal-ventral horn connectivity (dorsal: p < 0.001, ventral: p < 0.001); furthermore, the within-hemicord dorsal-ventral horn connectivity was significantly stronger than the between-hemicord dorsal-ventral horn connectivity (p = 0.001; but see results described under Aim 3).

### Aim 2 - segment-specific connectivity

Our second aim was to test whether resting-state connectivity could also be observed at the level of single spinal segments (Figure 4). Based on probabilistically defined spinal segments (Cadotte et al., 2015), it is evident that our acquired field of view contains the sixth, seventh and eighth cervical segments as well as the first thoracic segment (C6, C7, C8, T1).

**Figure 4:**
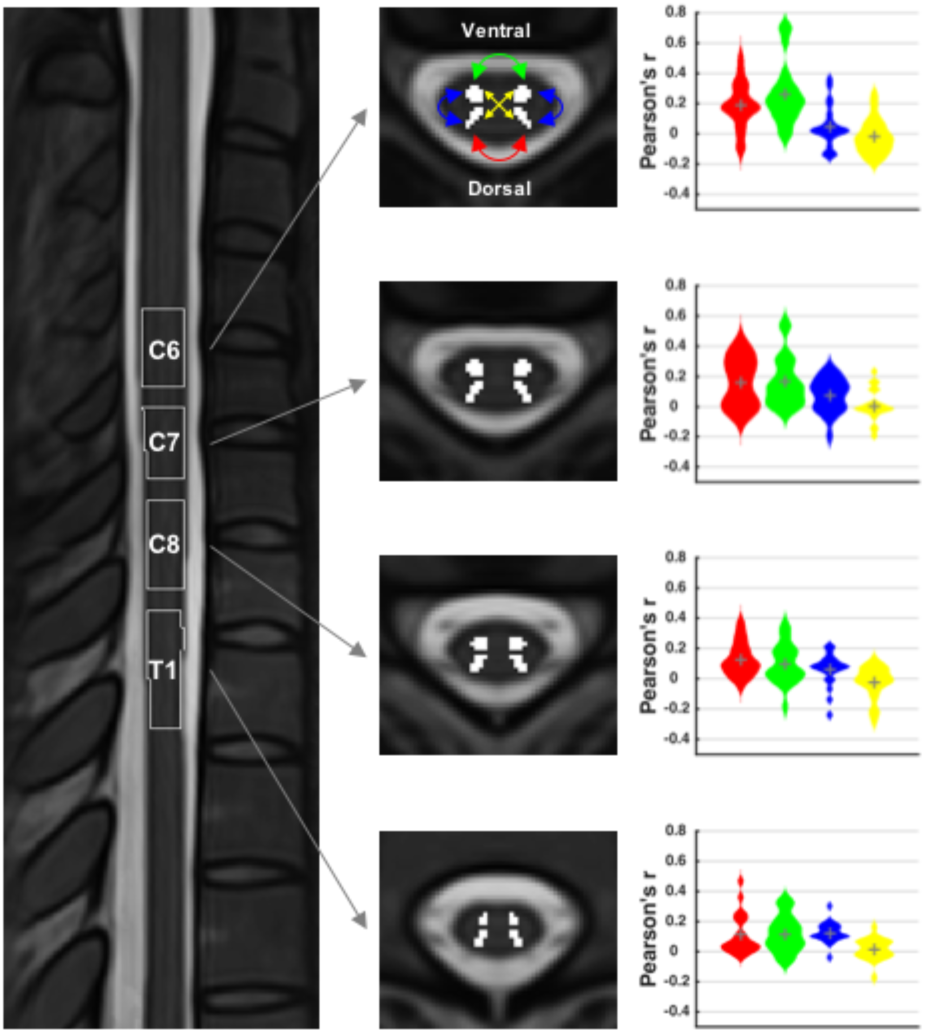
Segment-specific connectivity.

The image on the left is a midline sagittal slice through the T2-weighted MNI-Poly-AMU template, with the thresholded probabilistic segments overlaid as outlines. The four transversal slices in the middle are taken from the centre of each of the segments, with the four horn masks overlaid in white. The violin plots on the right demonstrate the correlation between the four horn masks within each segment as smoothed histograms of the distributions (the mean is indicated by the grey plus; color-coding as in Figure 3).

**Table 1.** Segment-specific connectivity. Shown are theresults for each of thefour spinal levels (C6,C7, C8, and T1) andeach of the fourcorrelations (betweendorsal horns, between ventral horns, between dorsal and ventralhorn within hemicords, betweendorsal and ventralhorn between hemicords), with *r* representing the average Pearson correlation, *p* representing the two-tailed family-wise-error corrected p-value from a permutation test, and % representing the percentage of participants showing a positive correlation.

|  | C6 | C7 | C8 | T1 |
| --- | --- | --- | --- | --- |
| **Dorsal-dorsal** |  |  |  |  |
| r | 0.19 | 0.16 | 0.13 | 0.11 |
| p | < 0.001 | 0.001 | < 0.001 | 0.005 |
| % | 85 | 80 | 95 | 85 |
| **Ventral-ventral** |  |  |  |  |
| r | 0.26 | 0.17 | 0.10 | 0.12 |
| p | < 0.001 | 0.001 | 0.01 | 0.001 |
| % | 95 | 85 | 75 | 80 |
| **Within-hemicord** |  |  |  |  |
| r | 0.05 | 0.08 | 0.06 | 0.12 |
| p | 0.35 | 0.03 | 0.11 | < 0.001 |
| % | 70 | 75 | 75 | 95 |
| **Between-hemicord** |  |  |  |  |
| r | -0.02 | 0.01 | -0.03 | 0.01 |
| p | 0.87 | 1 | 0.70 | 0.81 |
| % | 35 | 40 | 45 | 55 |

As can be seen in Table 1, the connectivity between dorsal horns as well as between ventral horns was highly significant at every single level, with a minimum of 80% of participants showing positive correlations between dorsal horns and a minimum of 75% of participants showing positive correlations between ventral horns at each spinal level (see also Figure 4). Connectivity between dorsal and ventral horns on the other hand was much more variable, with only some segments (C7 and T1) showing significant results for within-hemicord dorsal-ventral connectivity and none of the segments showing significant results for between-hemicord dorsal-ventral connectivity (Table 1 & Figure 4). These results thus corroborate the whole-cord connectivity results and show that this technique is indeed able to pick up relationships at the level of single spinal segments.

In a post-hoc analysis, we investigated whether dorsal and ventral horns in the different spinal segments would show a similar connectivity pattern. To this end, we correlated the connectivity pattern of each segment (i.e. the four obtained intra-segmental correlations) with every other segment within each participant and then averaged the results with respect to inter-segmental distance within each participant. This resulted in 20 (i.e. number of participants) correlations for i) one-segment distance, ii) two-segment distance, and iii) three segment distance. We then used two-tailed non-parametric permutation testing to investigate whether these correlations would be significantly positive or negative and indeed observed that they were all significantly positive, though becoming slightly weaker with distance (Figure 5): the correlation for one-segment distance was strongest (r = 0.40, t = 5.5, p_corrected_ < 0.001), followed by two-segment distance (r = 0.37, t = 4.8, p_corrected_ < 0.001) and three-segment distance (r = 0.22, t = 2.8, p_corrected_ = 0.03). This demonstrates that different spinal segments show similar ‘connectivity fingerprints’.

**Figure 5.**
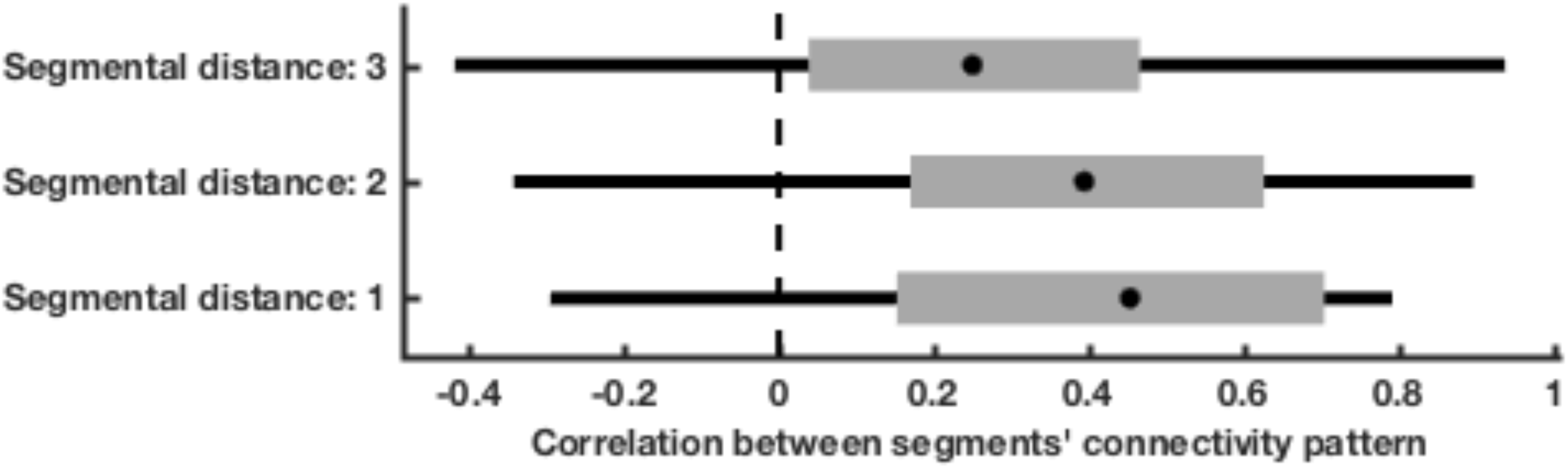
Similarity in segmental connectivity patterns.

The box-plots show how strongly the connectivity pattern (i.e. the four intra-segmental correlations) of each segment correlate with the connectivity pattern in every other segment, dependent on the distance between segments (one-segment distance: C6-C7, C7-C8, C8-T1; two-segment distance: C6-C8, C7-T1; three-segment distance: C6-T1). The filled black dot represents the group median, the edges of the boxes cover represent the 25^th^ and 75^th^ percentiles and whiskers encompass approximately 99% of the data.

### Aim 3 - robustness of connectivity

Our third aim was to assess how robust the observed resting-state connectivity would be against variations in the analysis pipeline, i.e. how reproducible the results would be across analyses. To this end, we carried out a total of 96 analyses using variations of ROI definition (two options), temporal filtering (two options), nuisance regression (eight options), and connectivity metric (three options); for brevity this was only done for the whole-cord connectivity.

First, we observed that the sign of the correlation (i.e. positive or negative connectivity) was remarkably robust against variations in the analysis pipeline for the dorsal horn correlations (96/96 analyses resulted in a positive group-average correlation), as well as for the ventral horn correlations (again, 96/96 analyses resulted in a positive group-average correlation). This was not the case for the correlations between ventral and dorsal horns, where within-hemicord correlations were at chance level (48/96 analyses resulted in a positive group-average correlation) and between-hemicord correlations were also rather variable (64/96 analyses resulted in a positive group-average correlation; Figure 6a). When investigating which analysis choices led to this variability, it became clear that for within-hemicord connectivity, ROI creation was the driving factor: positive connectivity was only observed with the PROB masks and negative connectivity was only observed with the COG masks. This suggests that time-course mixing due to the close proximity of the dorsal and ventral ROIs when using the PROB masks is the sole reason for observing positive correlations for within-hemicord dorsal-ventral connectivity, and that these are thus most likely artefactual. For between-hemicord dorsal-ventral connectivity, the influence of analysis choice was not so clear-cut and the only factor we could identify was the use of NS nuisance regression: negative correlations never occurred when this was employed.

**Figure 6.**
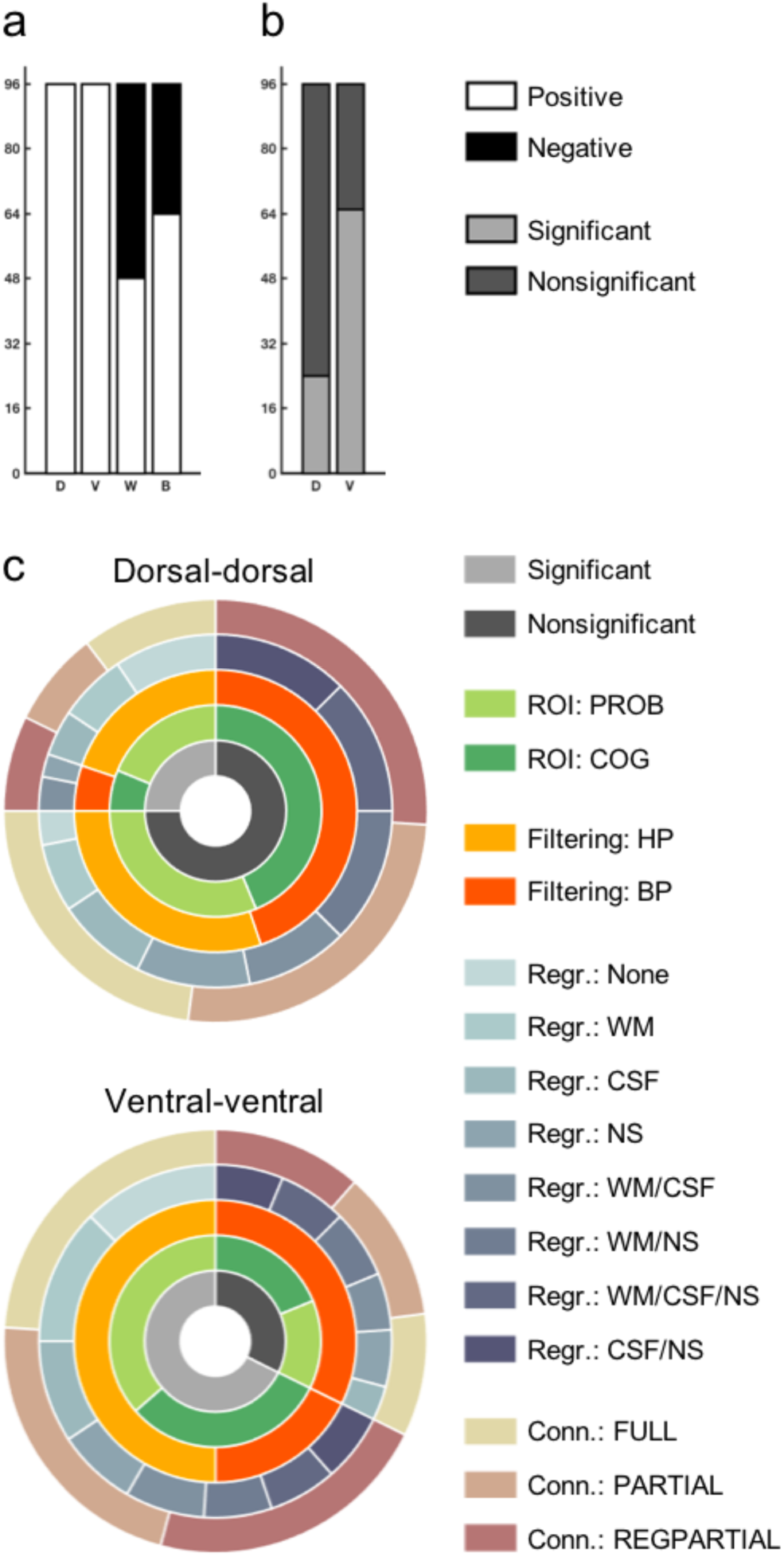
Robustness of connectivity.

**a)** Depicted are the number of analyses (out of a total of 96) that show positive (white) or negative (black) connectivity. Only dorsal-dorsal and ventral-ventral connectivity is robust against variations in the analysis pipeline. Abbreviations: D, dorsal-dorsal; V, ventral-ventral; W, within-hemicord dorsal-ventral; B, between-hemicord dorsal-ventral. **b)** Depicted are the number of analyses (out of a total of 96) that show significant (light grey) or nonsignificant (dark grey) connectivity, this time limited to dorsal-dorsal and ventral-ventral connectivity. Abbreviations: D, dorsal-dorsal; V, ventral-ventral. **c)** The radial bar-plots depict which analysis choices contribute to significant/nonsignificant connectivity as observed in **b)**. The innermost ring shows significance/nonsignificance and is a grouping factor for the next rings: ROI creation, temporal filtering, nuisance regression, and connectivity metric.

Considering that only connectivity between dorsal horns and connectivity between ventral horns seems to be stable across analysis choices, we limited our next analysis – where we assessed whether the significance of the correlations is influenced by variations in the data analysis pipeline – to these two correlations. For the connectivity between dorsal horns, only 24 out of 96 analyses showed a significant correlation, whereas for the connectivity between ventral horns, 65 out of 96 analyses showed a significant correlation (note though that this is a conservative estimate, as FWE-correction was based on four tests; Figure 6b). This worryingly low level of statistical robustness could be mainly explained by two factors (Figure 6c): temporal filtering and nuisance regression. For the ventral horns, 31 out of the 31 non-significant correlations could be explained by the use of band-pass filtering – within these 31 cases, different variations of nuisance regression were found to contribute as well, though no nuisance regression approach had as much of an impact as band-pass filtering. For the dorsal horns, 43 out of the 72 non-significant correlations could be explained by band-pass filtering – in contrast to the ventral horns nuisance regression had an even stronger impact here with a total of 69 of the 72 non-significant correlations due to variations in this approach, with NS nuisance regression having the strongest impact. Figure 6c shows how the levels of each factor (ROI creation, temporal filtering, nuisance regression, connectivity metric) are distributed over the significant and non-significant dorsal horn and ventral horn correlations.

Abbreviations: PROB, probabilistic masks; COG, centre of gravity masks; HP, high-pass temporal filtering; BP, band-pass temporal filtering; Regr., nuisance regression; WM, white matter; CSF, cerebrospinal fluid; NS, non-spinal; Conn., connectivity metric; FULL: full correlation; PARTIAL: partial correlation; REGPARTIAL, regularized partial correlation.

When using a repeated-measures ANOVA to investigate the effects of the different analysis choices on the correlation coefficients more generally (i.e. without binning the data into positive/negative or significant/non-significant), we made three main observations (Table 2): 1) nuisance regression had a consistently strong main effect on connectivity, 2) the main effects of temporal filtering and correlation metric were modest, and 3) ROI creation had a very large main effect on within-hemicord dorsal-ventral connectivity.

**Table 2.** Effects of variations in data analysis. Shown are the main effects of the four-way repeated-measures ANOVA that was carried out for each of the four horn-to-horn correlations and investigated the impact of different analysis choices. Please note that the subscript of each F-value represents the degrees of freedom of the corresponding test and that Greenhouse-Geisser correction was applied in case of non-sphericity (which can result in non-integer degrees of freedom).

| ROI creation | Temporal filtering | Nuisance regression | Connectivity metric |
| --- | --- | --- | --- |
| **Dorsal-dorsal** |  |  |  |
| $F_{1,19} = 3.0$ | $F_{1,19} = 0.0$ | $F_{7,133} = 13.7$ | $F_{1.2,23.3} = 6.8$ |
| p-value = 0.10 | p-value = 0.93 | p-value < 0.001 | p-value = 0.01 |
| **Ventral-ventral** |  |  |  |
| $F_{1,19} = 4.8$ | $F_{1,19} = 2.9$ | $F_{7,133} = 14.5$ | $F_{1.1,20.6} = 2.3$ |
| p-value = 0.04 | p-value = 0.10 | p-value < 0.001 | p-value = 0.15 |
| **Within-hemicord** |  |  |  |
| $F_{1,19} = 184.5$ | $F_{1,19} = 1.6$ | $F_{7,133} = 1.8$ | $F_{1.1,20.4} = 3.0$ |
| p-value < 0.001 | p-value = 0.22 | p-value = 0.10 | p-value = 0.10 |
| **Between-hemicord** |  |  |  |
| $F_{1,19} = 2.3$ | $F_{1,19} = 0.0$ | $F_{7,133} = 10.3$ | $F_{1.0,10.3} = 0.7$ |
| p-value = 0.15 | p-value = 0.88 | p-value < 0.001 | p-value = 0.42 |

Finally, we used two complementary inferential procedures (both of which make use of all 96 analyses) to test the robustness of the horn-to-horn connections. First, when testing whether the average of all 96 analyses was significant (after FWE-correction for four tests), we observed that the dorsal horn as well as the ventral horn connections were significant (dorsal-dorsal: p = 0.037, ventral-ventral: p = 0.001), whereas the dorsal-ventral connections were not (within-hemicord: p = 0.512, between hemicord: p = 1). Second, we used non-parametric combination (NPC) testing, and observed significant effects for the dorsal horn correlations (p = 0.013), the ventral horn correlations (p < 0.001) and the within-hemicord dorsal-ventral horn correlations (p = 0.017), but not for the between-hemicord dorsal-ventral horn correlations (p = 0.239).

## Discussion

In this resting-state fMRI study of the human spinal cord, we observed significant functional connectivity between the dorsal horns, as well as between the ventral horns, but not between the dorsal and ventral horns - neither within hemicords nor between hemicords. These effects were not only evident when considering the whole acquired extent of the cord, but also when considering data from single spinal segments. Finally, we could show that functional connectivity between the ventral horns and between the dorsal horns was mostly robust against variations in the data analysis pipeline, highlighting the inferential reproducibility of these effects.

The motivation for this study arose from results obtained by Barry and colleagues (2014, 2016), who – using ultra high field imaging (7T) of the spinal cord – demonstrated significant and reproducible resting-state functional connectivity between the dorsal horns as well as between the ventral horns. Considering that such connectivity measurements could be a useful tool for probing both disease progression and treatment response in neurological disorders with a strong spinal component, we aimed to test whether we could replicate these findings at the field strength of 3T which is much more prevalent in clinical settings. To this end we reanalysed a previously published resting-state data-set acquired at 3T (Kong et al., 2014) – in our previous publication we had used an exploratory ICA-based approach that is however not suitable for investigating connectivity between a-priori defined ROIs. Here we show that despite many technical differences between the study by Barry and colleagues and our study (e.g. field strength, hardware vendor, fMRI protocol, temporal degrees of freedom, cervical segments covered, estimation of connectivity) we can replicate their main findings at 3T: similar to them, we observed significant resting-state functional connectivity between the ventral horns (which are important for motor function), as well as between the dorsal horns (which are important for sensory function). Both of these findings were robust against inter-individual differences, with at least 90% of participants showing positive connectivity. Also mirroring Barry and colleagues’ findings, when averaging across the whole cord we did not observe significant correlations between the dorsal and ventral horns, neither when investigating within-hemicord connectivity nor when investigating between-hemicord connectivity. Consequently, when comparing connectivity strengths, we observed that both dorsal-dorsal and ventral-ventral connectivity was significantly stronger than dorsal-ventral connectivity, both within and between hemicords. In contrast to Barry et al. (2014) we did not observe a significant difference between ventral and dorsal connectivity (see also below). In any case, it is reassuring to see that largely similar results were obtained in these studies despite many technical differences – suggesting that resting-state spinal cord fMRI signals are a robust phenomenon, which bodes well for future multi-centre studies (such as currently underway for spinal cord diffusion tensor imaging: Samson et al., 2016).

Both Barry and colleagues’ and our findings were obtained when averaging data along the rostro-caudal axis of the spinal cord – in both instances data were averaged across the whole extent of the field of view. While this approach might help in removing noise and detecting functional connectivity patterns, it ignores the segmental structure of the spinal cord (Baron, 2015) and does not address whether it is possible to detect altered connectivity in localized cord regions as might occur for example with spinal cord compression (Nouri et al., 2015). We therefore tested whether we could detect resting-state connectivity within single spinal cord segments. Investigating this issue at the group level has only become possible very recently with the development of probabilistic maps of spinal segments (Cadotte et al., 2015) and their integration with a standard space template of the spinal cord (Fonov et al., 2014), both openly available with the Spinal Cord Toolbox (https://sourceforge.net/projects/spinalcordtoolbox; De Leener et al., 2016). With this approach we were able to demonstrate that the overall features of group-level resting-state connectivity – significant dorsal horn correlations and ventral horn correlations – were also detectable at every spinal level we investigated (sixth cervical to first thoracic level), though robustness against inter-individual differences was somewhat lower than for the whole-cord analysis. Segment-wise dorsal-ventral connectivity was never apparent between-hemicords and only partly within-hemicords, though this is most likely artefactual (see below). Interestingly, when investigating whether the pattern of intra-segmental connectivity remained similar across segments, we observed that this was indeed the case, though a trend for reduced similarity with increasing distance was apparent as well (suggesting that at large intersegmental distances there might indeed be different patterns of intra-segmental connectivity). At least for the segments we imaged, we can thus conclude that dorsal-dorsal and ventral-ventral connectivity seems to be a consistent feature, which our resting-state technique has the ability to detect and might thus be employed in studies investigating neurological disorders with localized spinal pathology.

Before spinal-cord resting-state signals can be used as potential biomarkers (Chen et al., 2015) it is important to assess how robust or reproducible they are (Barry et al., 2016). This has come to the forefront more generally in recent years with concerns regarding the reproducibility of published research in the biomedical literature and beyond (Begley and Ioannidis, 2015; Iqbal et al., 2016). We therefore set out to test how reproducible / robust our obtained results are against reasonable and common variations in the data-analysis pipeline. Note that we are not using the term “reproducibility” as it is used in computational science (where it refers to authors making raw data and analysis code available, so that others can recompute the original results; Peng, 2011), but rather in the form of robustness or inferential reproducibility (Goodman et al., 2016) – i.e. we aim to demonstrate that the basic inferences we draw here (there being strong ventral horn and dorsal horn connectivity) are not conditional on a specific analysis path and should thus be immune to biased data-analyses (Head et al., 2015; which have been discussed under terms such as “data-torturing” Mills, 1993; “researcher degrees of freedom” Simmons et al., 2011; and “p-hacking” Simonsohn et al., 2014); for the sake of brevity we limited these analyses to the whole-cord results.

While there is an enormous flexibility of fMRI data analysis pipelines (Carp, 2012), we chose to focus on a few analysis steps that have received attention in resting-state studies of the brain, namely ROI creation, temporal filtering, nuisance regression and connectivity metric (e.g. Pruim et al., 2015; Smith et al., 2011). We observed that the sign of the correlation was completely immune to changes in data analysis for dorsal horn connectivity as well as for ventral horn connectivity, but not for dorsal-ventral horn connectivity: within-hemicord dorsal-ventral connectivity only became positive when ROIs were used that were very close to each other, which likely led to time-course mixing (due to the spatial point-spread function of the BOLD response) and inflated positive correlations. We then focussed on the significance of dorsal horn connectivity and ventral horn connectivity and observed that in only about 25% of the different permutations of analysis approaches dorsal horn connectivity was significant; for the ventral horns this was higher with around 70%. When we investigated what was driving this worryingly low robustness, we observed that it was partly due to the use of band-pass filtering: when we ignored the analyses employing a band-pass filter, ventral horn connectivity was significant in 100% of the analyses and dorsal horn connectivity was significant in 40% of the analyses. This is interesting in light of recent findings in the spinal cord (Barry et al., 2016) and the brain (Pruim et al., 2015), where it was demonstrated that band-pass filtering had a negative impact on both the detectability and reproducibility of resting-state connectivity. Consistent with this notion, numerous brain imaging studies have shown that meaningful signal is also contained in higher frequencies above the traditional cut-off of 0.08Hz (Boubela et al., 2013; Chen and Glover, 2015; Gohel and Biswal, 2015; Niazy et al., 2011), which might hold true for the spinal cord as well (though thorough characterizations using short-TR data are needed). In addition to these descriptive results, we used two significance tests based on data-aggregation across all 96 analyses and observed significant effects for both dorsal-dorsal and ventral-ventral connectivity in both of these tests, supporting the robustness of these connections. Further support for the idea that dorsal-dorsal and ventral-ventral connectivity is not artificially induced by common noise sources is provided by the fact that full correlation and partial correlation analyses showed similar results.

When investigating the effects of analysis variation more generally using a repeated measures ANOVA based on the correlation coefficients, we observed that nuisance regression had the most consistent influence, while the influence of connectivity metric was rather weak (in line with the above-mentioned findings, temporal filtering most strongly affected ventral horn correlations and ROI creation had its strongest effect on within-hemicord dorsal-ventral correlations). It will be important to tease apart the influence of the various nuisance regression approaches, especially when considering that dorsal horn connectivity seems to be quite prone to these analysis-specific influences. In our opinion the reason for this susceptibility to nuisance regression choice is most likely related to the shape and location of the dorsal horns in comparison to the ventral horns: 1) the dorsal horns are more elongated and thinner, making them more susceptible to partial volume effects with white matter and 2) they border the posterior edge of the cord and could thus be susceptible to partial volume effects with CSF. It remains to be seen how one can obtain the best balance between preserving “true” signal and removing noise, the latter of which is especially important for resting-state data to avoid false positives due to noise-driven spurious correlations (Cole et al., 2010; Murphy et al., 2013) and even more so in the spinal cord due to its higher level of physiological noise compared to the brain (Cohen-Adad et al., 2010; Piché et al., 2009).

Aside from these technical considerations, an obvious question pertains to the neurobiological underpinnings of the observed signals. This has been discussed in detail previously (e.g. with regard to possible influences of central pattern generators on ventral correlations, input from the peripheral nervous system on dorsal correlations, and supra-spinal influences on both of these; see Barry et al., 2014; Eippert and Tracey, 2014; Kong et al., 2014) and we will only briefly discuss possible underlying factors not mentioned previously. With regard to the dorsal horn connectivity, there is some anatomical evidence for primary afferents that also cross to the contralateral side (Culberson et al., 1979; Light and Perl, 1979) and electrophysiological evidence for an interneuronal network that connects the two dorsal horns (Fitzgerald, 1982, 1983). This has been corroborated more recently with a number of studies identifying populations of dorsal commissural interneurons (Petkó and Antal, 2000; Bannatyne et al., 2006), though these mostly focussed on the lumbar spinal cord. With regard to the ventral horns, there is a wealth of literature on commissural interneurons (for review, see e.g. Jankowska, 2008). While most of these investigations occurred in the upper cervical segments or in the lumbar cord, there is also evidence for commissural systems in the lower cervical segments that we investigated (Alstermark and Kümmel, 1990; Soteropoulos et al., 2013). One should also not discount the effect of respiration on the observed ventral horn resting-state connectivity. Respiration is typically treated as a source of physiological noise in spinal fMRI, e.g. due to breathing-induced B_0_ shifts (Verma and Cohen-Adad, 2014) and breathing-induced modulation of CSF flow (Schroth and Klose, 1992b). However, the activity of respiratory motoneurons (Lane, 2011; Monteau and Hilaire, 1991) in the ventral horns might actually underlie some of the observed ventral horn resting-state connectivity. While motoneurons innervating the primary expiratory muscles are unlikely to play a role (the abdominal and internal intercostal muscles are generally not recruited during quiet breathing and are furthermore innervated only from thoracic and lumbar segments), the main inspiratory muscle – the diaphragm – is innervated via the phrenic nerve which originates from segments C3 to C5 in humans (Hollinshead and Keswani, 1956; Routal and Pal, 1999). In this regard it is interesting to note that Barry et al. (2014, 2016) – who acquired data from these segments – observed much stronger correlations between the ventral horns than the dorsal horns, whereas we – who acquired data from below these segments – did not observe such a pronounced difference. Also, resting-state connectivity between ventral horns was almost non-existent in monkeys who were mechanically ventilated during anaesthesia (Chen et al., 2015). Such an interpretation could be tested by investigating whether ventral horn connectivity changes during breathing manipulations. Furthermore, even the ventral horn connectivity we observed could be driven by respiratory factors, because respiratory interneurons as well as respiratory motoneurons innervating the scalene muscles are present in the lower cervical spinal cord (Lane, 2011; Monteau and Hilaire, 1991; see also Wei et al., 2010). The contribution from these neurons is somewhat unclear however, as the function of respiratory interneurons during normal breathing remains to be elucidated and the scalenes are generally considered only accessory respiratory muscles (but see De Troyer and Estenne, 1984), which also show much lower discharge rates than the diaphragm (Saboisky et al., 2007).

Whatever the neurobiological underpinnings of the observed resting-state signals are, it is important to point out several limitations of the present report. *First*, we need to acknowledge that the observed correlations are rather weak (group-average r of less than 0.3 in most cases), which could stem from the lower temporal signal-to-noise ratio (tSNR) of spinal fMRI data due to the inherent limitations in data acquisition from this structure. While we employed an fMRI protocol that was optimized to minimize CSF inflow effects and susceptibility-induced signal drop-out (Finsterbusch et al., 2012), we cannot rule out that these factors still had a detrimental effect on ventral and dorsal horn signals. It is also worth mentioning that we did not spatially smooth the data: while smoothing will boost the tSNR, it would also introduce a large amount of time-course mixing between the ROIs and was thus omitted. In the future one might consider using more advanced approaches such as smoothing solely along the cord axis (De Leener et al., 2016) or non-local spatial filtering (Manjón et al., 2010). It is also possible that our denoising approach was not successful in characterising and removing noise sources properly, but we think this to be unlikely considering the variety of (mostly validated) methods we have employed for noise removal. A final – and neurobiologically more interesting - consideration relates to the possibility that the resting-state signal in each horn of a segment is not only determined by inputs from other horns of this segment, but might be strongly influenced by inter-segmental input (see also Kong et al., 2014) as well as input from supraspinal regions and the peripheral nervous system. This relates to a *second* point, namely that we only investigated intra-segmental connectivity in detail, but not inter-segmental connectivity, as this would have far exceeded the scope of this report. *Third*, it is currently unclear why we were not able to obtain evidence for dorsal-ventral connectivity when considering that both within-hemicord and between-hemicord connectivity is essential for some sensorimotor functions such as reflexes – while absence of evidence obviously does not imply evidence of absence, it might just be the case that spinal cord resting-state fluctuations do not cycle through the whole anatomical repertoire of connections and that tonic inhibition of such a system might play a role. *Fourth*, it is important to point out that with the chosen spatial resolution (1x1x5mm), all ROI time-series will be subject to a certain amount of time-course mixing between grey matter and white matter due to partial volume effects. And *fifth*, while we investigated the issue of reproducibility or robustness against variations in data analysis, it will also be crucially important for future studies to investigate the inter-session reliability of spinal cord resting-state signals over days, weeks and months, before they might be used in clinical settings (Zuo and Xing, 2014). Acquiring several resting-state sessions in the same participants over time (for first steps in this direction at 3T, see San Emeterio Nateras et al., 2016 and Liu et al., 2016) would also allow to determine which of the methods we have employed here is optimal in terms of providing the least variable results across sessions.

## Conclusions

In this study we have replicated previously obtained 7T resting-state results (Barry et al., 2014) at the more widely available field strength of 3T. We have furthermore shown that ROI-based dorsal horn connectivity as well as ventral horn connectivity was highly significant not only when averaged across the length of the cord, but also at each of the acquired spinal cord segments, which exhibited similar connectivity patterns. Finally, we have investigated the issue of robustness/reproducibility and observed that the obtained results are mostly robust against variations in data analysis. In our opinion this suggests that functional connectivity might be a methodologically robust tool for investigating basic spinal cord research questions, such as the correspondence between resting-state and task-based connectivity (Cole et al., 2014) in the spinal cord, the integration between spinal and supra-spinal processes in health (Büchel et al., 2014) and disease (Freund et al., 2016), or how tonic pain protocols (Segerdahl et al., 2015) might lead to a change and possibly spread of dorsal horn spontaneous fluctuations. Even more important, this technique could complement current approaches for assessing pathology, disease progression, and treatment response in neurological disorders with a profound spinal cord component, such as spinal cord injury.

## Acknowledgements

AMW received support from the National Research Council of Brazil (CNPq; 211534/2013-7). CB is supported by the DFG (FOR 1328 and SFB 936) and ERC (2010-AdG_20100407). IT is supported by a Wellcome Trust Strategic Award and the NIHR Oxford Biomedical Research Centre. JCWB was supported by the Medical Research Council UK (G0700238). JLA is supported by a Wellcome Trust Strategic Award (098369/Z/12/Z).

